# Impact of cross-ancestry genetic architecture on GWAS in admixed populations

**DOI:** 10.1101/2023.01.20.524946

**Authors:** Rachel Mester, Kangcheng Hou, Yi Ding, Gillian Meeks, Kathryn S. Burch, Arjun Bhattacharya, Brenna M. Henn, Bogdan Pasaniuc

## Abstract

Genome-wide association studies (GWAS) have identified thousands of variants for disease risk. These studies have predominantly been conducted in individuals of European ancestries, which raises questions about their transferability to individuals of other ancestries. Of particular interest are admixed populations, usually defined as populations with recent ancestry from two or more continental sources. Admixed genomes contain segments of distinct ancestries that vary in composition across individuals in the population, allowing for the same allele to induce risk for disease on different ancestral backgrounds. This mosaicism raises unique challenges for GWAS in admixed populations, such as the need to correctly adjust for population stratification to balance type I error with statistical power. In this work we quantify the impact of differences in estimated allelic effect sizes for risk variants between ancestry backgrounds on association statistics. Specifically, while the possibility of estimated allelic effect-size heterogeneity by ancestry (HetLanc) can be modeled when performing GWAS in admixed populations, the extent of HetLanc needed to overcome the penalty from an additional degree of freedom in the association statistic has not been thoroughly quantified. Using extensive simulations of admixed genotypes and phenotypes we find that modeling HetLanc in its absence reduces statistical power by up to 72%. This finding is especially pronounced in the presence of allele frequency differentiation. We replicate simulation results using 4,327 African-European admixed genomes from the UK Biobank for 12 traits to find that for most significant SNPs HetLanc is not large enough for GWAS to benefit from modeling heterogeneity.

## Introduction

The success of genomics in disease studies depends on our ability to incorporate diverse populations into large-scale genome-wide association studies (GWAS)^1–4^. Cohort and biobank studies are growing to reflect this diversity^5–7^, and a variety of techniques exist which incorporate populations of different continental ancestries into GWAS^8^. However, while admixture has been an important factor in other steps in the disease mapping process, such as fine-mapping^9^ and estimating heritability^10,11^, individuals of mixed ancestries (admixed individuals) have largely been left out of traditional association studies. GWAS performed in admixed populations have greater power for discovery compared to similar sized GWAS in homogeneous populations^12,13^. Thus, excluding admixed individuals from association studies will not only increase health disparities, but will also disadvantage other populations. To prevent this exclusion, approaches to association studies have been developed specifically for admixed populations^14–17^. However, the impact of HetLanc (differences in estimated allelic effect sizes for risk variants between ancestry backgrounds) on GWAS methods remains underexplored. Of particular interest are recently admixed populations, defined as less than 20 generations of mixture between two ancestrally distinct populations. In such populations, the admixture process creates mosaic genomes comprised of chromosomal segments originating from each of the ancestral populations (i.e., local ancestry segments). Local ancestry segments are much larger than linkage disequilibrium (LD) blocks^18^; thus, LD patterns within each local ancestry block of an admixed genome reflect LD patterns of the ancestral population. Similarly, allele frequency estimates from segments of a particular local ancestry are expected to reflect allele frequencies of the ancestral population. Variation in local ancestry across the genome leads to variability in global ancestry (the average of all local ancestries within a given individual). Such variability in local and global ancestries could pose a problem to GWAS in admixed populations as genetic ancestries are often correlated with socio-economic factors that also impact disease risk, thus yielding false positives in studies that do not properly correct for genetic ancestries. Because local and global ancestry are only weakly correlated^19^, complete control of confounding due to admixture requires conditioning on both local and global ancestry^20^. However, the success of admixture mapping indicates that the possibility of losing power due to over-correction for local ancestry stratification is serious^17,21^.

GWAS in admixed populations is typically performed either using a statistical test that ignores local ancestry altogether (e.g., the Armitage trend test, ATT) or using a test that explicitly allows for HetLanc (e.g., Tractor). The former provides superior power in the absence of HetLanc with the latter having great potential for discovery in its presence. However, these methods’ relative statistical power for discovery depends on the cross-ancestry genetic architecture of the trait: i.e., which variants are causal and what are those variants’ ancestry-specific frequencies, causal effects, and linkage disequilibrium patterns. For example, existing studies have found that ATT can yield a 25% increase in power over Tractor^3^ in the absence of HetLanc while Tractor has higher power when causal effects are different by more than 60%^15^. However, the full impact of cross-ancestry genetic architecture on GWAS power in admixed populations remains under-explored.

In this work, we use simulations to perform a comprehensive evaluation quantifying the impact of these factors on the power of GWAS approaches in admixed populations. We provide guidelines for when to use each test as a function of cross-ancestry genetic architecture. Elements of cross-ancestry genetic architecture such as allele frequencies, global ancestry ratios, and LD are known or can be calculated in advance of a GWAS to determine which of our simulation results apply in each case. Using extensive simulations, we find that ATT should be preferred when HetLanc is small or non-existent. We quantify the extent of HetLanc and the ancestry-specific allele frequency differences required for Tractor to overcome the extra degree of freedom penalty. We further validate our results using the African-European admixed population in the UK Biobank (UKBB). By examining the HetLanc of significant SNPs in the UKBB, we can understand how often it rises to a level that impacts the power of traditional GWAS.

## Results

### Heterogeneity by Local Ancestry Impacts Association Statistics in Admixed Populations

HetLanc occurs when a SNP exhibits different estimated allelic effect sizes depending on its local ancestry background. HetLanc can manifest itself at causal SNPs due to genetic interactions between multiple causal variants or differential environments, although recent work suggests that the magnitude and frequency of these types of epistatic effects between causal variants is limited^22^. A more common form of HetLanc is observed at non-causal SNPs that tag the causal effect in a differential manner across ancestries. Differential linkage disequilibrium by local ancestry at these non-causal SNPs (tagged SNPs) can cause HetLanc even when allele frequencies and causal effect sizes are the same across ancestries. The extent to which HetLanc exists and the magnitude of these differences in effect sizes are yet uncertain^22–38^. However, the existence of HetLanc plays an important role in the power of GWAS methods to detect associations. Consider the example in Figure 1 in which the allelic effect size for a tagged SNP is estimated for a phenotype in an admixed population. In this population, both the tagged SNP and the true causal SNP may exist in regions attributed to both local ancestries present in the population (Figure 1a). Since LD patterns differ by local ancestry, the correlation between the tagged and causal SNPs will also depend on local ancestry (Figure 1b). This differential correlation between tagged and causal SNPs will cause the estimated allelic effect size for the tagged SNP 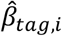 to depend on local ancestry *i* (Figure 1c). Thus, even for cases in which true causal effect sizes are the same across ancestries, allelic effect sizes estimated for the tagged SNP may be heterogeneous. Since GWAS cannot determine true causal effect sizes, we introduce *R_het_*, a measure of HetLanc which allows for both true causal effect-size heterogeneity and LD- and allele frequency-induced estimated allelic effect-size heterogeneity.

**Figure 1:**
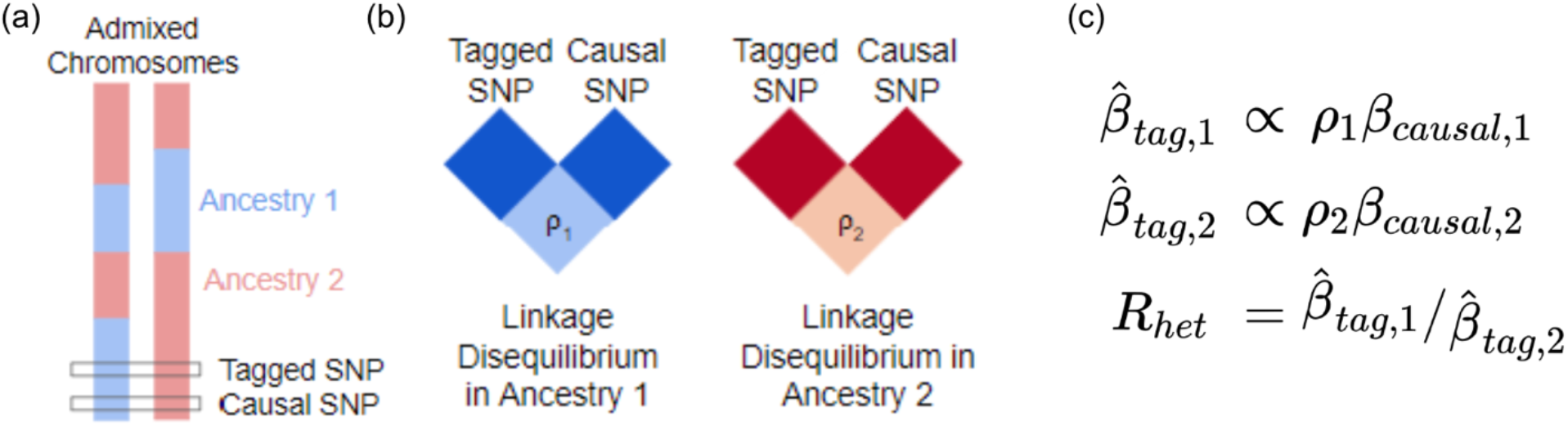
Toy example of how differential LD by local ancestry can induce HetLanc. **(a)** Admixed populations contain haplotypes with different local ancestry at the causal or tagged SNP. **(b)** The correlation between tagged and causal SNPs depends on their local ancestry due to differential LD by local ancestry. **(c)** In a GWAS, the estimated marginal SNP effect size is proportional to the true causal effect size and the correlation between the tagged and causal SNPs 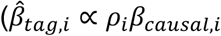, where *i* refers to the *i_th_* ancestry).

### Methods for association testing in admixed populations

We start with a formal definition for a full model relating genotype, phenotype, and ancestry for a single causal SNP:

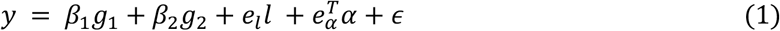

where *y* is a phenotype, *g*_1_, and *g*_2_ are vectors that represent the number of alternate alleles with local ancestry 1 and 2 (such that *g*_1_, + *g*_2_ = *g*, the genotype in standard form), *β*_1_, and *β*_2_ are ancestry-specific marginal effect sizes of the SNP, *l* is the vector of local ancestry counts at the locus, *e_l_* is the effect size of *l, α* is a matrix of other covariates such as global ancestry, 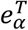 is the vector of the effect sizes of *α*, and *ϵ* is random environmental noise.

Variability across local and global ancestries has been leveraged in various statistical approaches for disease mapping in admixed populations. One of the first methods developed for association was admixture mapping (ADM)^17,29^. ADM tests for association between local ancestry and disease status in cases and controls or in a case-only fashion. This association is achieved by contrasting local ancestry deviation with expectations from per-individual global ancestry proportions. Therefore, ADM is often underpowered especially in situations in which allele frequency at the causal variant is similar across ancestral populations^30^. Genotype association testing is traditionally performed using an Armitage trend test (ATT). ATT tests for association between genotypes and disease status while correcting for global ancestry to account for stratification^17,31^. However, neither ADM nor ATT take advantage of the full disease association signal in admixed individuals. SNP1, SUM, and MIX are examples of association tests that combine local ancestry and genotype information. SNP1 regresses out local ancestry in addition to global ancestry to control for fine-scale population structure. This approach helps control for fine-scale population stratification but may remove the signal contained in local ancestry information^32^. SUM^33^ combines the SNP1^14^ and ADM statistics into a 2 degree of freedom test. MIX^14^ is a case-control test that incorporates SNP and local ancestry information into a single degree of freedom test. Most recently Tractor^15^ conditions the effect size of each SNP on its local ancestry followed by a joint test allowing for different effects on different ancestral backgrounds. This step builds the possibility of HetLanc explicitly into the model, which may be particularly important when SNPs are negatively correlated across ancestries^34^. Other varieties of tests have also been developed using different types of frameworks, most notably BMIX^34^ which leverages a Bayesian approach to reduce multiple testing burden. These statistics have been compared at length^3,14,17,35^. However, existing comparisons do not consider HetLanc, nor do they thoroughly discuss allele frequency differences across ancestries.

### ATT has more power than Tractor in the absence of heterogeneity by ancestry

First, we use simulations to compare type I error and power for each association statistic in Table 1. Starting with 10,000 simulated admixed individuals based on a 50/50 admixture proportion, we simulate 1,000 case-control phenotypes with a single causal SNP (see Methods). We calculate type I error as the probability of each method to detect significant associations in non-causal SNPs (see Methods). Type I error is well controlled for every association test, well under the 5% threshold expected by the chosen p-value (Figure 2a). The mean type I error was ≤ 4.36 × 10^−2^% for every association test. The maximum value was ≤ 0.6% for every association test. We next calculate power to detect SNPs with an odds ratio of *OR*_1_, = *OR*_2_ = 1.2 (see Methods). We find that SNP1 had the highest power at 42.14%. However, SNP1 was not significantly more powerful than either MIX (power 42.12%, p-value 0.878) or ATT (power 42.05%, p-value 0.325, Figure 2b). The power of all three of these tests was significantly higher (p-value ≤ 1 × 10^−16^) than for SUM (power = 33.44%), ADM (power = 0.039%), or Tractor (power = 31.89%). Thus, we find that while these association statistics are all well controlled, power does substantially differ between them. In the absence of both HetLanc and allele frequency difference, 1 degree of freedom SNP association tests outperform 2 degree of freedom tests.

**Table 1:** Summary of GWAS association statistics. All tests adjust for global ancestry and can be used on binary traits, and all tests except MIX can be implemented with adjustment for additional covariates and use on quantitative traits. For more information on the comparison of ATT, ADM, SUM, and MIX see^32^. We note that while additional methods exist^35–39^ we do not focus on them in this work because they do not directly relate to equation 1.

| Association<br>Statistic | Statistical Test<br>($H_0$) | Assumptions<br>on $\beta$ | Covariates | Degrees of<br>Freedom |
| --- | --- | --- | --- | --- |
| ADM | $e_l = 0$ | -- | $\alpha$ | 1 |
| ATT | $\beta = 0$ | $\beta = \beta_1 = \beta_2$ | $\alpha$ | 1 |
| SNP1 | $\beta = 0$ | $\beta = \beta_1 = \beta_2$ | $l, \alpha$ | 1 |
| MIX | $e_l \circ \beta = 0$ | $\beta = \beta_1 = \beta_2$ | $\alpha$ | 1 |
| SUM | $\beta = 0$ and $e_l = 0$ | $\beta = \beta_1 = \beta_2$ | $l, \alpha$ | 2 |
| Tractor | $\beta_1 = 0$ and $\beta_2 = 0$ | -- | $l, \alpha$ | 2 |

**Figure 2:**
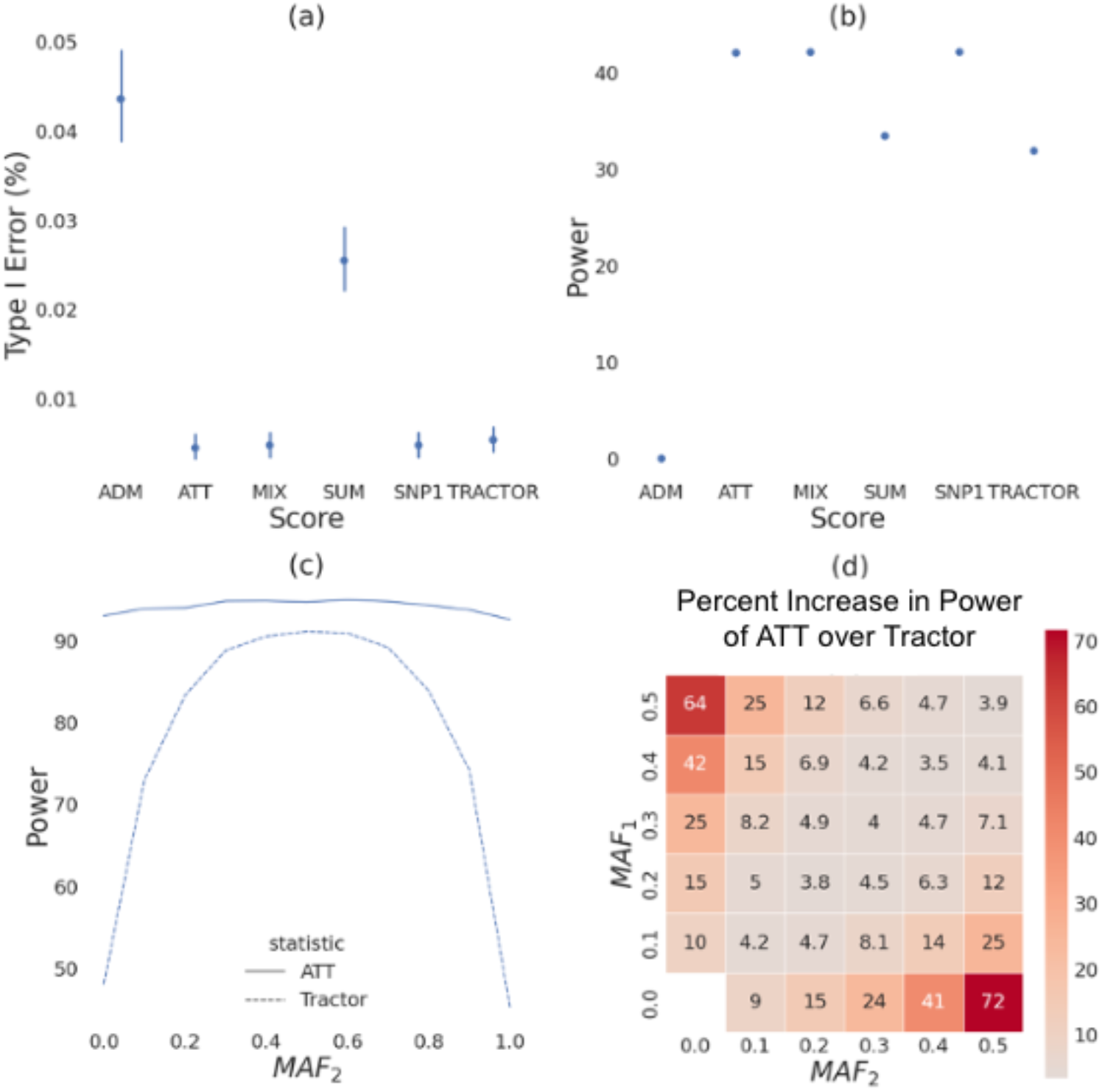
Association statistics in the absence of HetLanc. **(a)** Type I error for association statistics. Type I error calculated as the probability for detecting significant association for a null SNP. 95% confidence interval shown. **(b)** Power for association statistics. Power calculated at odds ratios *OR*_1_, = *OR*_2_ = 1.2. 95% confidence interval too narrow for display. **(c)** Power for ATT and Tractor as *MAF*_2_ is varied between 0.0 and 1.0 and *MAF*_1_, is fixed at 0.5. Power for both methods varies as MAF difference varies. 95% confidence interval too narrow for display. **(d)** Heatmap of percent increase in power of ATT over Tractor when *β*_1_, = *β*_2_ = 1.0. Minor allele frequencies *MAF*_1_, and *MAF*_2_ varied from 0.0 to 0.5 in increments of 0.1. All simulations are for a case-control **(a-b)** or quantitative **(c-d)** traits simulated 1,000 times for a population of 10,000 individuals with global ancestry proportion 50/50. Power calculated using a Bonferroni-corrected threshold of standard threshold p-value < 1 × 10^−5^ **(a-b)** or standard threshold p-value <5 ×10^−8^ **(c-d)**. Case-control traits **(a-b)** have case-control ratio 1:1 and 10% case prevalence, quantitative traits **(c-d)** have heritability *h*^2^ = 0.005. Heritability, global ancestry, causal effect size *β* and overall MAF do not qualitatively impact these results (Figures S1, S2 and S3).

We next investigate how differences in minor allele frequency (MAF) impact the power of ATT and Tractor in the case where true causal effect sizes are the same. We investigate the impact of varying MAF in each ancestry independently. Using our 10,000 simulated admixed individuals from the previous experiment, we simulate 1,000 quantitative phenotypes with a single causal SNP (see Methods). First, we let *MAF*_1_, = 0.5 and *MAF*_2_ range from 0.0 to 1.0 with a 0.1 increment and plot power over *MAF*_2_ (Figure 2c). We find that ATT has higher power than Tractor at all levels of MAF difference. Since Tractor has an extra degree of freedom compared to ATT, Tractor is disadvantaged when *β*_1_, = *β*_2_. When *MAF*_1_, = *MAF*_2_, ATT has 94.7% power, with Tractor at 91.1% power. However, as *MAF*_2_ becomes more different from *MAF*_1_, ATT maintains its power at 93.0%. By contrast, Tractor loses much of its power, with only 45.3% power when the causal allele is fixed at 100% in population 2 and only 48.1% power when the causal allele is absent in population 2. ATT maintains higher power than Tractor even at varying levels of heritability (Figures S1, S2, S3), *MAF*_1_, (Figure S1), global ancestry (Figure S2), and effect size *β* (Figure S3). However, the difference in power has a large range depending on the MAF difference between local ancestries.

Next we introduce percent difference in power, a one-dimensional metric to compare between these association statistics (see Methods). We use this metric to visualize how varying MAF_1_ and MAF_2_ independently impacts the power of ATT and Tractor (Figure 2d). The percent increase in power when using ATT over Tractor when the causal SNP is absent in population 2 is 64%. The power difference between ATT and Tractor increases as MAF difference increases. Furthermore, the lower the MAF starts out in population 1, the larger the power difference between these two statistics. Specifically, when *MAF*_1_, = 0.5 and *MAF*_2_ = 0.1, the difference in MAF is 0.4 and ATT has a 25% power increase over Tractor. However, when *MAF*_1_, = 0.4 and *MAF*_2_ = 0.0, the difference in MAF is still 0.4 but ATT has a 42% increase in power over Tractor.

While this result corroborates previous studies^40–42^, the relationship between Tractor and admixture mapping provides insight into the mechanism behind this dynamic. Mainly, as allele frequency differentiation by local ancestry increases, so does the power of the admixture mapping test statistic. In fact, ADM has no power when minor allele frequencies do not differ by ancestry but achieves up to 6.7% power when MAF_1_ = 0.0 and MAF_2_ = 0.5 (Figure S4a). However, the Tractor method uses the admixture mapping statistic as its null hypothesis. A stronger null hypothesis will be rejected less often than a weaker one even when the alternative hypothesis is the same, causing any test utilizing a strong null hypothesis to have less power. Thus, Tractor will have less power when its null hypothesis (ADM) has more power, which occurs in situations with high allele frequency differentiation. When allele frequencies do not differ by ancestry, Tractor achieves 91% power in our simulations. However, when MAF_1_ = 0.0 and MAF_2_ = 0.5 Tractor power plummets to 44% (Figure S4b).

While high levels of allele frequency differentiation drastically decrease the power of Tractor, ATT also has a smaller decrease in power at high levels of allele frequency differentiation, from 95% at equal allele frequencies to 93% when MAF_1_ = 0.0 and MAF_2_ = 0.5 (Figure S4c). This decrease in power is not as large as that suffered by Tractor, but it is also due to increased power of the null hypothesis at higher frequency differentiation across populations. The null hypothesis of the ATT test statistic only includes global ancestry, but the power of global ancestry alone to predict a trait increases as allele frequency differentiation increases^32^. The idea that including global ancestry as a covariate in these analyses reduces power for SNPs with large MAF differences raises the question of how much attenuation can be expected when more exact measures of global ancestry (such as principal components) are included in the analysis. However, the overall power attenuation due to the inclusion of global ancestry is small compared to that due to local ancestry; thus, we shift our focus back to considering local ancestry-specific effects on power.

### Impact of HetLanc on Power Depends on Allele Frequency Differences

Next, we investigate the impact of MAF differences and HetLanc on power differences between ATT and Tractor. The exact relationship between HetLanc (measured as R_het_), MAF difference, and percent difference in power is complex (Figure 3a). First, there is a window when 0.5 < *R_het_* < 1.5 in which, regardless of MAF difference, HetLanc is not enough to empower Tractor over ATT. Thus, at these “low” levels of HetLanc, ATT will reliably have more power than Tractor across the allele frequency spectrum. Similarly, when *R_het_* < −0.5, there is no allele frequency difference which would empower ATT over Tractor. This corroborates our findings that when effect sizes are in opposite directions, Tractor is expected to have improved power over ATT regardless of MAF difference. We can see that it is characteristics of both ATT and Tractor that drive this trend (Figure S8). The power of ATT depends most strongly on the magnitude of R_het_ and is diminished the most when effect sizes are in opposite directions. By contrast, the power of Tractor depends strongly on both MAF difference and R_het_. These two factors combine to create an asymmetric shape for the percent difference in power (Figure 3a). This asymmetry in power observed for the Tractor method is likely due to correlations between effective sample size, allele frequency, global ancestry, and local ancestry that can occur in an asymmetric manner when causal effect sizes and minor allele frequencies differ between local ancestries^32^. We additionally investigate similar scenarios with varied global ancestry proportions (Figure S5), heritability (Figure S6), and population-level MAF (Figure S7). While the exact boundaries of these regions do differ, the overall shape of this heatmap and the conclusions mentioned above do not qualitatively change.

**Figure 3:**
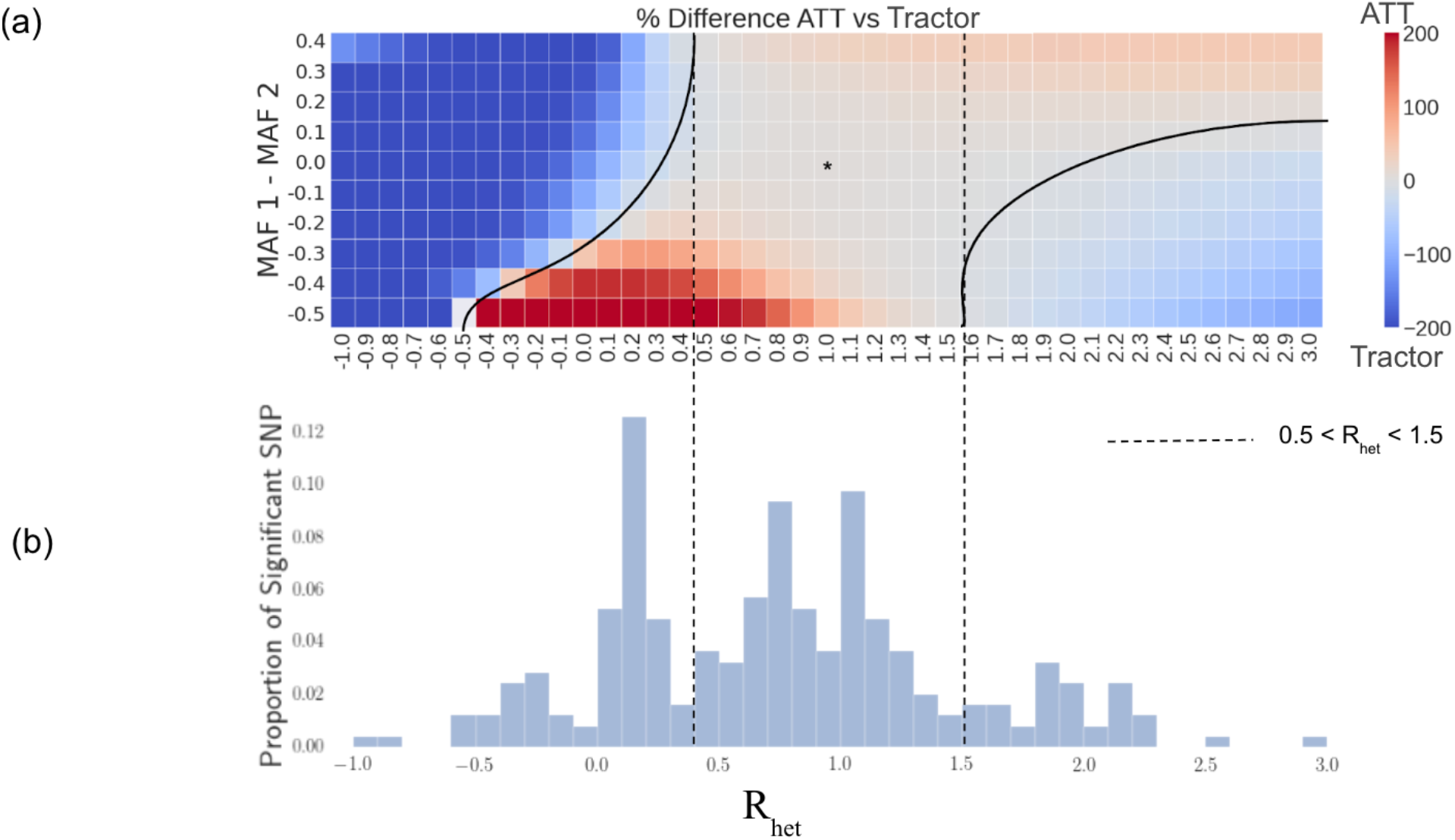
Impact of HetLanc on percent difference in power depends on MAF difference. **(a)** Heatmap of percent difference in power for ATT vs Tractor. The “*” indicates the center with no HetLanc or MAF difference. Quantitative trait simulated 1,000 times for a population of 10,000 individuals on a trait with effect size *β*_1_, ranging from −1.0 to 3.0 in increments of 0.1, and effect size *β*_2_ = 1.0. Global ancestry proportion 50/50, heritability at *h*^2^ = 0.005, and minor allele frequencies *MAF*_1_, = 0.5 and *MAF*_2_ ranging from 0.1 to 1.0 in increments of 0.1. Power calculated using a standard threshold p-value < 5 × 10^−8^. **(b)** Histogram of empirical *R_het_* 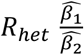 for significant SNPs found for 12 phenotypes in the UKBB. 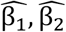 estimated using Tractor.

### ATT Finds More Significant Loci Across 12 Traits in the UK Biobank

We next seek to understand the impact of correcting for local ancestry in genetic analyses in real data. We investigate both Tractor and ATT in individuals with African-European admixture in the UK Biobank. These individuals have on average 58.9% African and 41.1% European ancestry over the population of 4,327 individuals. First, we investigate MAF differences between segments of African and European local ancestry over 16,584,433 imputed SNPs. We find that 72.8% of them have an absolute allele frequency difference of < 0.115 across local ancestry (Figure S10).

Next, we investigate empirically derived values of R_het_ to determine in which region of the heatmap estimated effect sizes are likely to be found in real data (Figure 3b). We ran the Tractor method on 12 quantitative traits to find the actual values of Rhet for the estimated effect sizes *β_AFR_* and *β_EUR_*. These traits were aspartate transferase enzyme (AST), BMI, cholesterol, erythrocyte count, HDL, height, LDL, leukocyte count, lymphocyte count, monocyte count, platelet count, and triglycerides. Then, we line up the histogram of these empirically derived values of *R_het_* with the heatmap. We find that for 69.3% of all significant SNPs, the empirical value for *R_het_* is within this [−0.5, 1.5] window. While this is an estimate, we predict the true difference between estimated marginal effect sizes might be smaller than indicated by these empirical values because Tractor is more powerful in identifying SNPs with heterogenous effect sizes. This result reflects previous findings that causal effects are similar across ancestries within admixed populations^22^. Due to this similarity in effect size, most of the significant SNPs sit in the center of the heatmap. This region of this heatmap predicts ATT will have more power than Tractor. We can compare the mean adjusted chi-square statistic of the SNPs found to be significant in this case. We find that this statistic is significantly larger for the ATT method than the Tractor method (Figure S9). For significant SNPs, the mean ATT 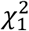 is 42.9, the mean adjusted Tractor 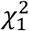 is 37.5, and the p-value for the difference is 2.11 × 10^−4^.

In addition to assessing HetLanc directly, we can also compare the number of independent significant SNPs found by ATT and Tractor for these phenotypes. We find that while the number of independent significant SNPs varies across all traits (Table S1), overall ATT finds more significant independent signals than Tractor (Figure 4a). We find 22 independent significant loci, with 19 loci found in ATT and 10 found in Tractor. This trend is most pronounced in HDL, in which 5 independent loci were determined to be significant by ATT compared to none for Tractor. Similarly, BMI, leukocyte count, and monocyte count also only had independent significant loci when testing using ATT as opposed to Tractor. Cholesterol and LDL had significant loci found by both ATT and Tractor, with a larger number found by ATT. Height is the only trait for which Tractor identified one significant locus but not ATT. Unfortunately, our sample sizes were not large enough to detect any significant loci for platelet count, triglycerides, or lymphocyte count. All significant loci for these 12 phenotypes are detailed in Table S1.

**Figure 4:**
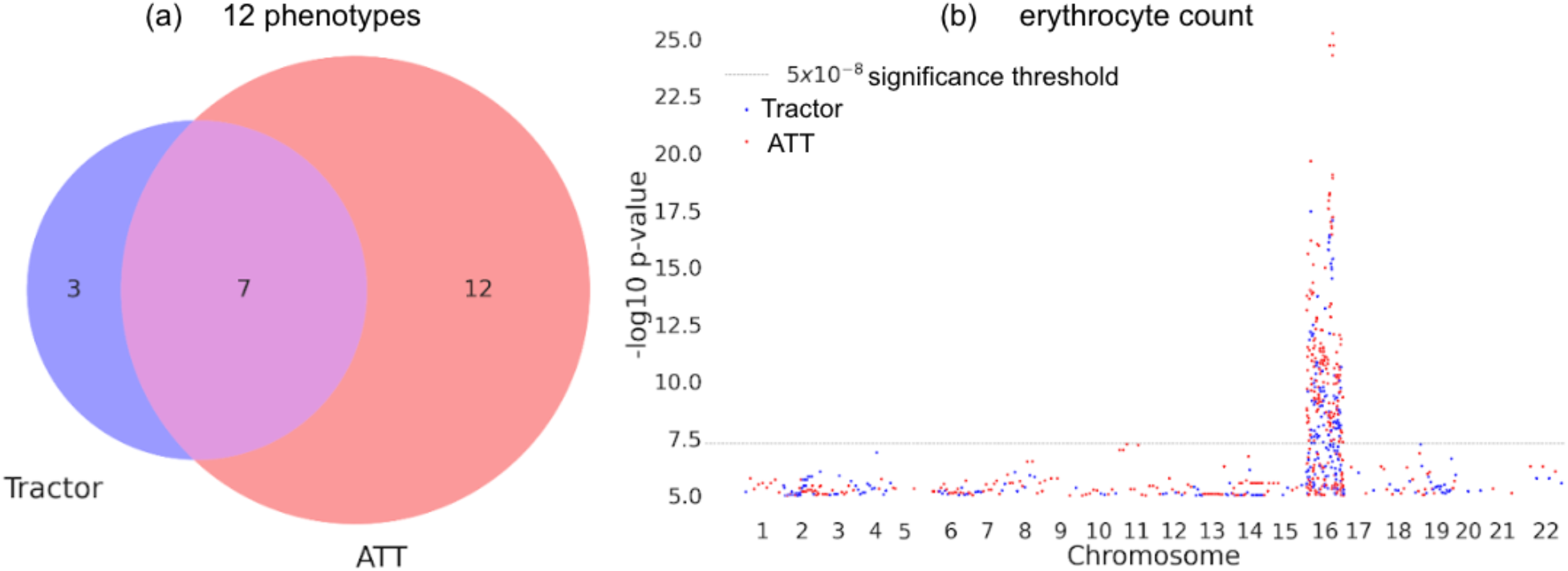
Comparing significant SNPs found with ATT and Tractor. **(a)** Venn diagram of independent significant loci found using ATT and Tractor in the UKBB across 12 quantitative traits. **(b)** Manhattan plot of chromosome for erythrocyte count in the UKBB. Significant SNPs found with ATT shown in red and significant SNPs found with Tractor shown in blue.

Additionally, we find that while ATT often finds more significant independent loci than Tractor, the two methods do not always find the same loci. Erythrocyte count is one phenotype in which we find an equal number of independent significant loci using both ATT and Tractor. However, not all loci overlap. Investigating the Manhattan plot of erythrocyte count specifically (Figure 4b) we see that loci on chromosome 16 are found by both ATT and Tractor. But outside of the main locus, both ATT and Tractor find separate additional significant regions. At the main locus, this Manhattan plot clearly shows that ATT has significantly smaller p-values for the same locus. Thus, in a smaller sample size only ATT would have found this important region. This example highlights the importance of choosing the most highly powered association statistic for any given situation. Manhattan plots for other phenotypes can be found in Figure S11.

## Discussion

In this work, we seek to understand the impact that estimated allelic effect-size heterogeneity by ancestry (HetLanc) has on the power of GWAS in admixed populations. Our main goal is to find whether conditioning disease mapping on local ancestry leads to an increase or decrease in power. We find that HetLanc and MAF differences are the two most important factors when considering various methods for disease mapping in admixed populations. We consider two association statistics - ATT, which ignores local ancestry, and Tractor, which conditions effect sizes on local ancestry. We find that in cases with small or absent levels of HetLanc, ATT is more powerful than Tractor in simulations of quantitative traits. This conclusion holds across a variety of global ancestry proportions and SNP heritabilities. We find that as MAF differentiation between ancestries increases, so does the improvement of power of ATT compared to Tractor. At high HetLanc (R_het_ > 1.5) or when effect sizes are in opposing directions (R_het_ < −0.5), we find that Tractor out-performs ATT. For African-European admixed individuals in the UKBB, most significant loci have both small measured HetLanc and MAF differences. We find that across 12 quantitative traits, ATT finds more significant independent loci than Tractor. Furthermore, ATT has smaller p-values for the loci that it shares with Tractor. This suggests that on smaller datasets more of the shared loci would be found by ATT than by Tractor.

This work has several implications for GWAS in admixed populations. Our results suggest that usually, ATT adjusted for global ancestry is the most powerful way to perform GWAS in an admixed population. However, it may be possible to predict the comparative power of ATT and Tractor using the allele frequencies and linkage disequilibria of a specific sample. Additionally, since in real analyses ATT and Tractor often find different loci, it is important to keep both methods in mind when performing analyses. These methods prioritize different types of loci, with ATT likely prioritizing loci with higher MAF differences and Tractor prioritizing loci with higher levels of HetLanc. From both scientific and social perspectives, it is important that admixed populations are incorporated more effectively into genetic studies. By providing insight into the strengths and limitations of these methods, we hope to enable studies to maximize their power in admixed populations.

We conclude with caveats and limitations of our work. When hoping to understand these patterns of power for association statistics, there are many combinations of different elements of genetic architecture to consider. These include phenotypic factors such as environmental variance and polygenicity, as well as elements of admixture such as the number of generations of admixture and the strength of linkage disequilibrium. We could not consider them all, and thus it is likely that additional nuances to our findings exist when other factors are considered. One major element not considered in this work is case-control traits. While we chose to focus on quantitative traits in this analysis due to their importance in simplicity and ubiquity, case-control traits are also important in medicine. It is possible that the behavior of these phenotypes will vary compared to the quantitative traits that we analyze here, both in simulations and real data. We suggest case-control traits as an interesting avenue of research for future works. Lastly, we chose to focus our analyses on ATT and Tractor due to their popularity and ease of use. We compare how these methods work “out of the box” to provide simple and usable guidance for others. However, as discussed in the introduction to this work, a variety of other association tests exist. It is likely that in certain circumstances one of these existing methods would outperform both ATT and Tractor.

## Acknowledgements

The authors would like to acknowledge Ella Petter, Ruth Johnson, and Vidhya Venkateswaran for their insightful feedback. RM supported in part by National Institutes for Health (NIH) award no. T32HG002536 and BMH and GLM supported in part by NIH grant R35GM133531 to BMH. The content is solely the responsibility of the authors and does not necessarily represent the official views of the NIH.

## Data and Code Availability

Code for this project, including simulation experiments, data processing pipeline, are available at https://github.com/rachelmester/AdmixedAssociation. An application for UK Biobank individual-level genotype and phenotype data can be made at http://www.ukbiobank.ac.uk.

## Methods

### Simulated Genotypes and Phenotypes

We simulate genotypes using the following procedure:

1. Draw global ancestry proportions *α* ~ *N*(*θ, σ*^2^) for 10,000 individuals where *θ* is the expected global ancestry proportion (either 0.5, 0.6, or 0.8) of ancestry 2, and *σ*^2^ is the variance of global ancestry in the population (*σ*^2^ = 0.125). We use *σ* ^2^ = 0.125 to reflect the variance of global ancestry found in the UK Biobank admixed population. *α* is coerced to 0 if it is negative and 1 if it’s larger than 1.
2. For each individual, draw a local ancestry count *l* ~ *Binomial*(*α*, 2), where *l* represents the local ancestry count of ancestry 2.
3. For each local ancestry, draw a genotype *g_i_* ~ *Binomial*(*l, f_i_*), where *f_i_* represents the minor allele frequency at local ancestry *i*.

We simulate phenotypes using the following procedure:

1. Standardize genotypes so that they have a mean 0 and variance 1.
2. Given some effect sizes *β*_1_, *β*_2_ calculate *Var_g_* = (*β*_1_*g*_1_ + *β*_2_*g*_2_)^2^, where *Var_g_* is the genetic variance component of the phenotypes.
3. Given some heritability *h*^2^, calculate 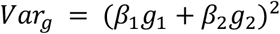, where *Var_e_* is the environmental variance component of the phenotypes. This comes from the equation 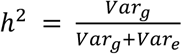.
4. For each individual, draw *ϵ* ~ *N*(0, *Var_e_*) where *ϵ* is the random noise to add to the phenotype to represent environmental variables.
5. Repeat for 1,000 replicates.

### Real Genotypes and Phenotypes

For our real data analysis, we used genotypes from the UK Biobank. We limited our study to participants with admixed African-European ancestry. Overall, we had 4,327 individuals with an average of 58.9% African and 41.1% European ancestry. We used the imputed genotypes for these individuals with a total of 16,584,433 SNPs. We calculated the top 10 PCs for these genotypes and added these PCs as covariates to all analyses as our global ancestry component. The phenotypes we used are also from the UK Biobank, and include aspartate transferase enzyme (AST), BMI, cholesterol, erythrocyte count, HDL, height, LDL, leukocyte count, lymphocyte count, monocyte count, platelet count, and triglycerides. We log transformed AST, BMI, HDL, leukocyte count, lymphocyte count, monocyte count, platelet count, and triglycerides to analyze all 12 traits as quantitative, continuous traits. We standardized all genotypes and phenotypes to be mean centered at 0.0 and have a standardized variance of 1.

### Association Testing

#### Simulated Data

We calculate the ATT and Tractor association tests on simulated data using scripts that can be found on https://github.com/rachelmester/AdmixedAssociation. ATT is a 1 degree of freedom association test that uses the model 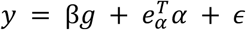 to test for *β* = 0 against a null hypothesis that includes global ancestry (*α*). Tractor is a two degree of freedom association test that uses the model 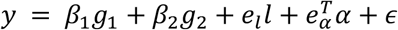 to test for *β*_1_, = 0 and *β*_2_ = 0 against a null hypothesis that includes local ancestry (*l*) and global ancestry (*α*). They can both be adapted to be used on case-control phenotypes or to adjust for additional covariates such as age and sex. For our simulations, we used global ancestry proportions as our measure of global ancestry (*α*). and did not need to adjust for any additional covariates such as age and sex as we did not model those factors in our simulations. For power calculations, we use a standard significance threshold of p-value < 5 × 10^−8^.

#### Real Data

We used admix-kit (https://kangchenghou.github.io/admix-kit/index.html) to perform the ATT and Tractor association tests on this data and extracted the p-values. In order to determine significant SNPs, we filtered for SNPs with a standard p-value of <5 × 10^−8^. For the Manhattan plots, we plot all SNPs with a p-value < × 10^−3^ for computational plotting purposes. For the Venn diagrams, in order to determine whether SNPs were part of the same loci, we grouped SNPs within a 500kB radius, and kept the most significant SNP from each test (ATT and Tractor) in that locus.

### Measures Used to Compare Our Results

In this work, we introduce several key measures that we use to compare our results. The formal definitions of these are the following:

<u>Percent difference in power:</u> 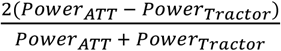
<u>Adjusted chi square:</u> We take the p-value from a *χ*^2^ statistic and convert it back to a 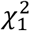 statistic, regardless of the original degrees of freedom. The adjusted chi square score for a 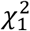 is itself.

