## Supplemental Material for "Impact of cross-ancestry genetic architecture on GWAS in admixed populations"

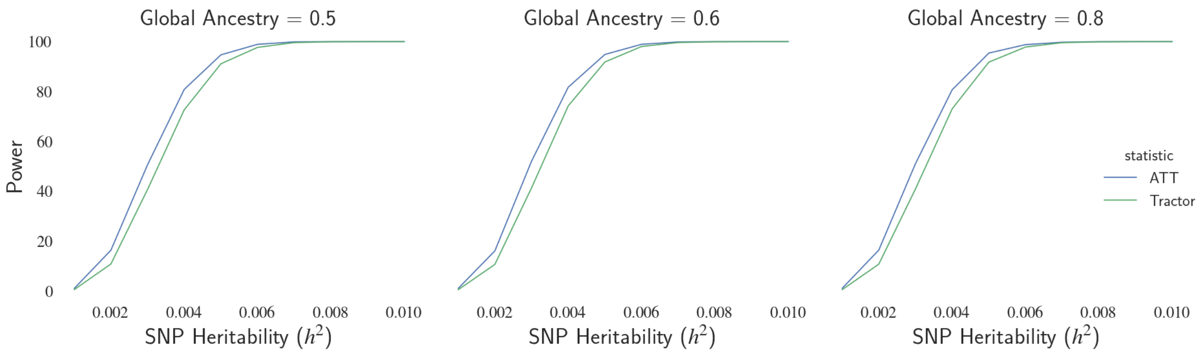


**Figure S1: Global ancestry does not have a large impact on power compared to choice of test statistic and SNP heritability.** Power curves of ATT and Tractor as SNP heritability varies. In this case where neither frequency nor causal effect size vary by local ancestry, ATT has increased power over Tractor, especially at small SNP heritabilities. Simulation results of 1,000 replicates with N = 10,000 individuals with minor allele frequency $MAF_1 = MAF_2 = 0.5$, and causal effect sizes $\beta_1 = \beta_2 = 1.0$. 95% confidence interval too narrow for display.

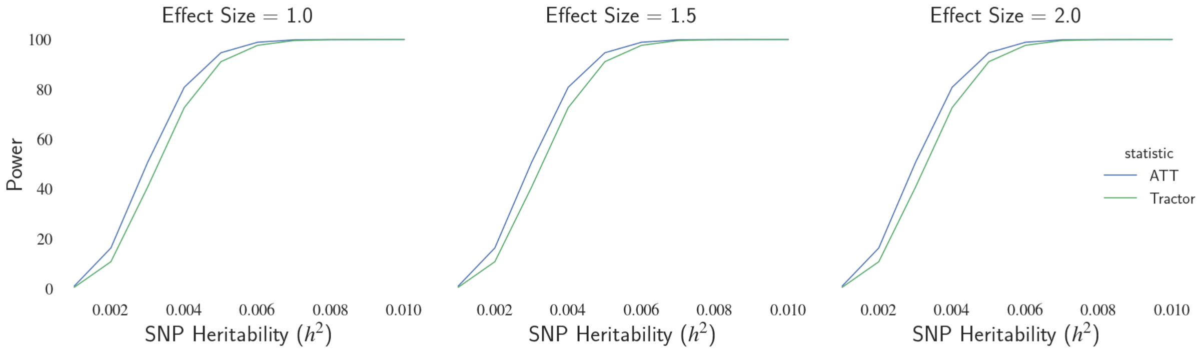


**Figure S2: Effect size does not have a large impact on power compared to choice of test statistic and SNP heritability.** Power curves of ATT and Tractor as SNP heritability varies. In this case where neither frequency nor causal effect size vary by local ancestry, ATT has increased power over Tractor, especially at small SNP heritabilities. Simulation results of 1,000 replicates with N = 10,000 individuals with minor allele frequency $MAF_1 = MAF_2 = 0.5$, global ancestry proportions at 50/50, and causal effect sizes $\beta_1 = \beta_2$. 95% confidence interval too narrow for display.

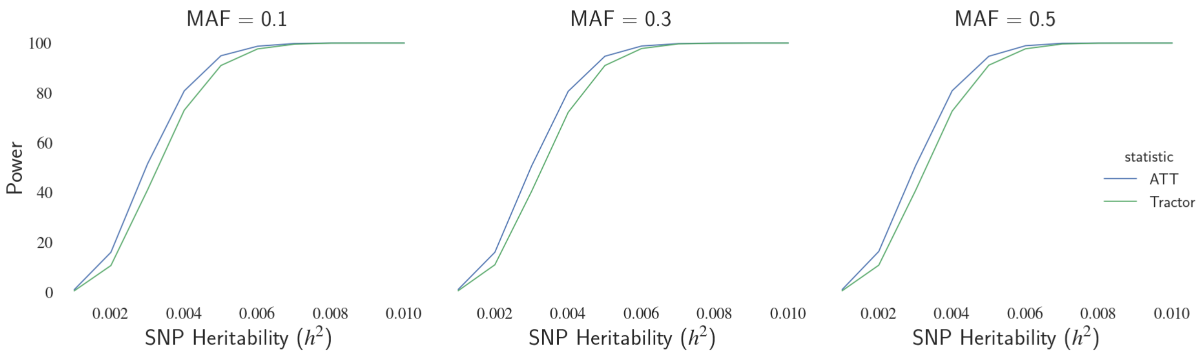


**Figure S3: Minor allele frequency does not have a large impact on power compared to choice of test statistic and SNP heritability.** Power curves of ATT and Tractor as SNP heritability varies. In this case where neither frequency nor causal effect size vary by local ancestry, ATT has increased power over Tractor, especially at small SNP heritabilities. Simulation results of 1,000 replicates with N = 10,000 individuals with minor allele frequency $MAF_1 = MAF_2$, global ancestry proportions at 50/50, SNP heritability $h^2 = 0.005$, and causal effect sizes $\beta_1 = \beta_2 = 1.0$. 95% confidence interval too narrow for display.

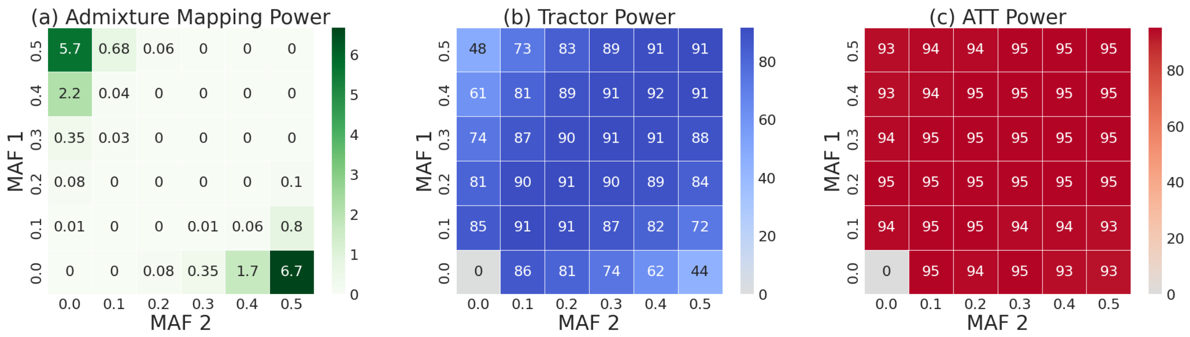


**Figure S4: Association statistic power at differing levels of allele frequency difference. (a)** Admixture mapping has maximum power when minor allele frequency difference by local ancestry is increased. **(b)** Tractor has drastically decreased power when minor allele frequency difference by local ancestry is increased. In this case where causal effect size does not vary by local ancestry, the decrease in Tractor power at high levels of minor allele frequency difference by local ancestry is driven by the increase in power for admixture mapping, which serves as the null hypothesis against which Tractor tests SNP-level effects. **(c)** ATT has slightly decreased power when minor allele frequency difference by local ancestry is increased. ATT does not suffer from using ADM as its null hypothesis as Tractor does, but the decrease in power is likely due to increased correlation between global and local ancestry at high levels of allele frequency difference. All panels are simulation results of 1,000 replicates with N = 10,000 individuals with global ancestry proportions at 50/50, SNP heritability $h^2 = 0.005$, and causal effect sizes $\beta_1 = \beta_2 = 1.0$.

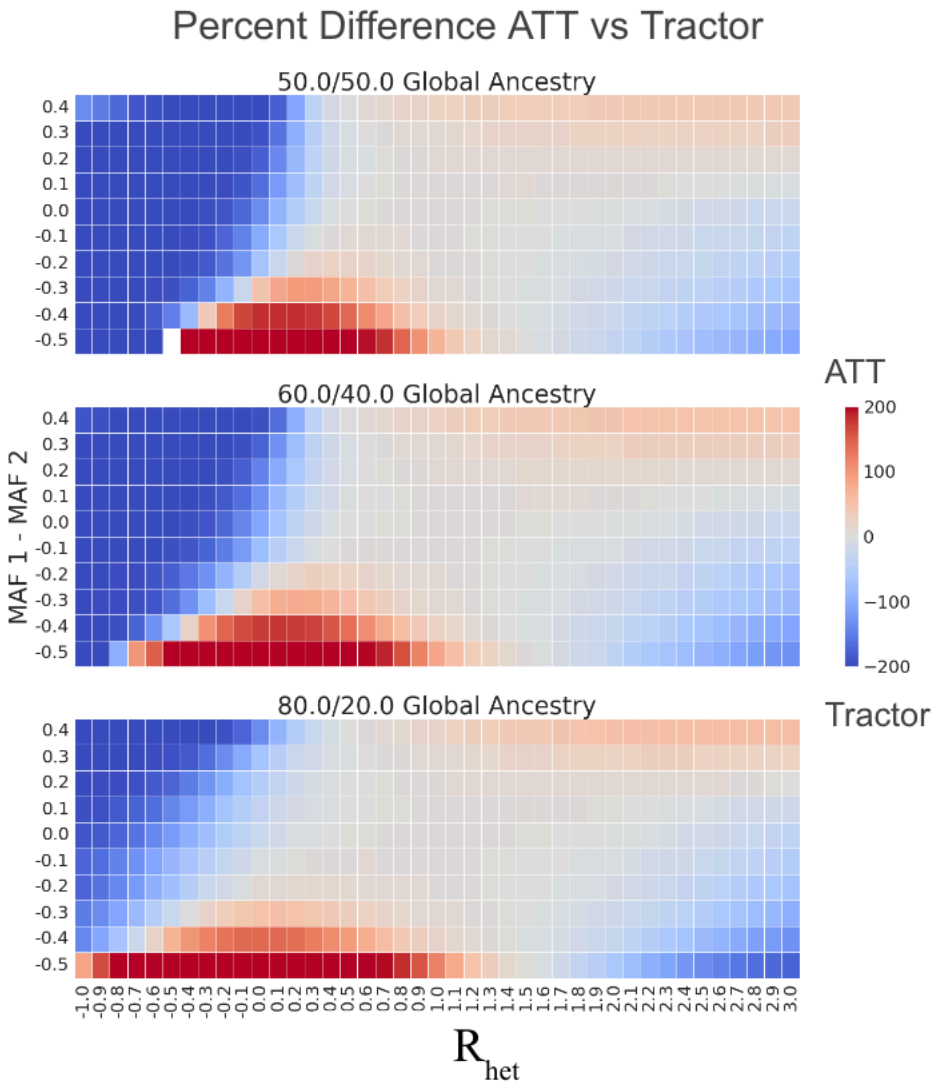


**Figure S5: Impact of HetLanc and MAF difference on percent difference in power depends on global ancestry ratios**. Heatmap of percent difference in power for ATT vs Tractor. Red indicates where $Power_{ATT} > Power_{Tractor}$. As global ancestry ratios become further from 50%, the range of HetLanc and MAF difference in which ATT has more power than Tractor increases. Simulation results of 1,000 replicates with N = 10,000 individuals with minor allele frequency $MAF_1 = 0.5$, heritability $h^2 = 0.005$, and causal effect size $\beta_2 = 1.0$.

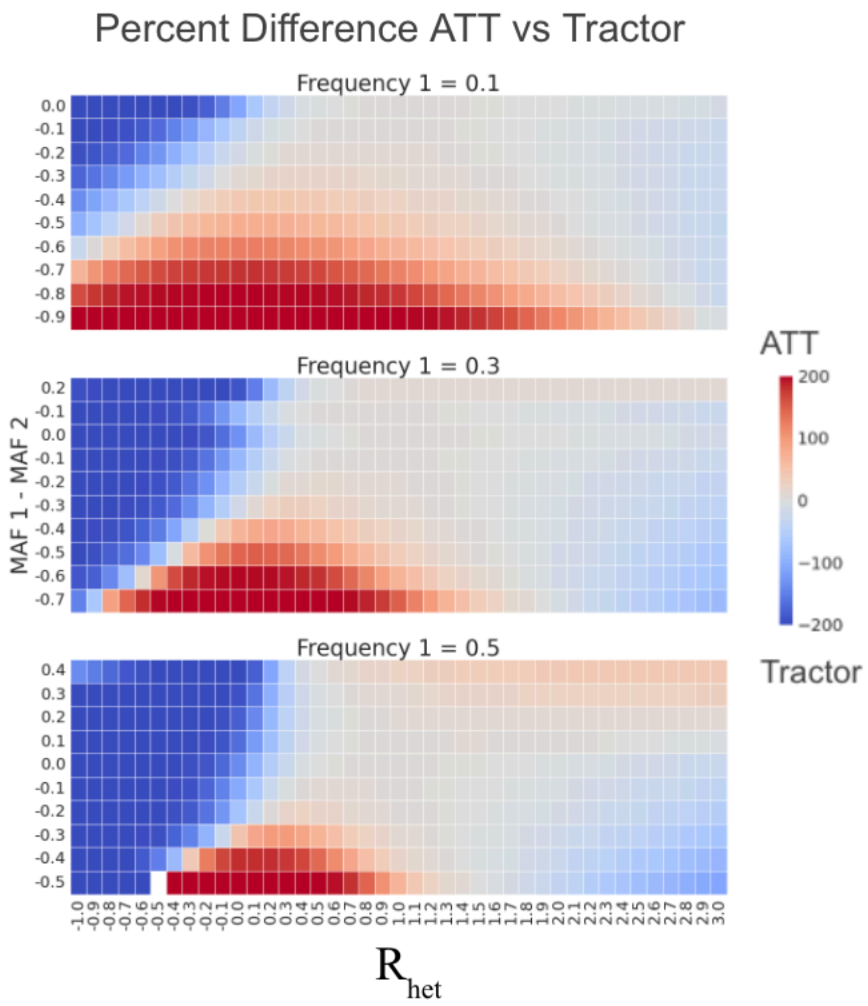


**Figure S6: Impact of HetLanc and MAF difference on percent difference in power depends on MAF.** Heatmap of percent difference in power for ATT vs Tractor. Red indicates where $Power_{ATT} > Power_{Tractor}$. As MAF becomes further from 0.5, the range of HetLanc and MAF difference in which ATT has more power than Tractor increases. Simulation results of 1,000 replicates with N = 10,000 individuals with global ancestry proportions at 50/50, SNP heritability $h^2 = 0.005$, and causal effect size $\beta_2 = 1.0$.

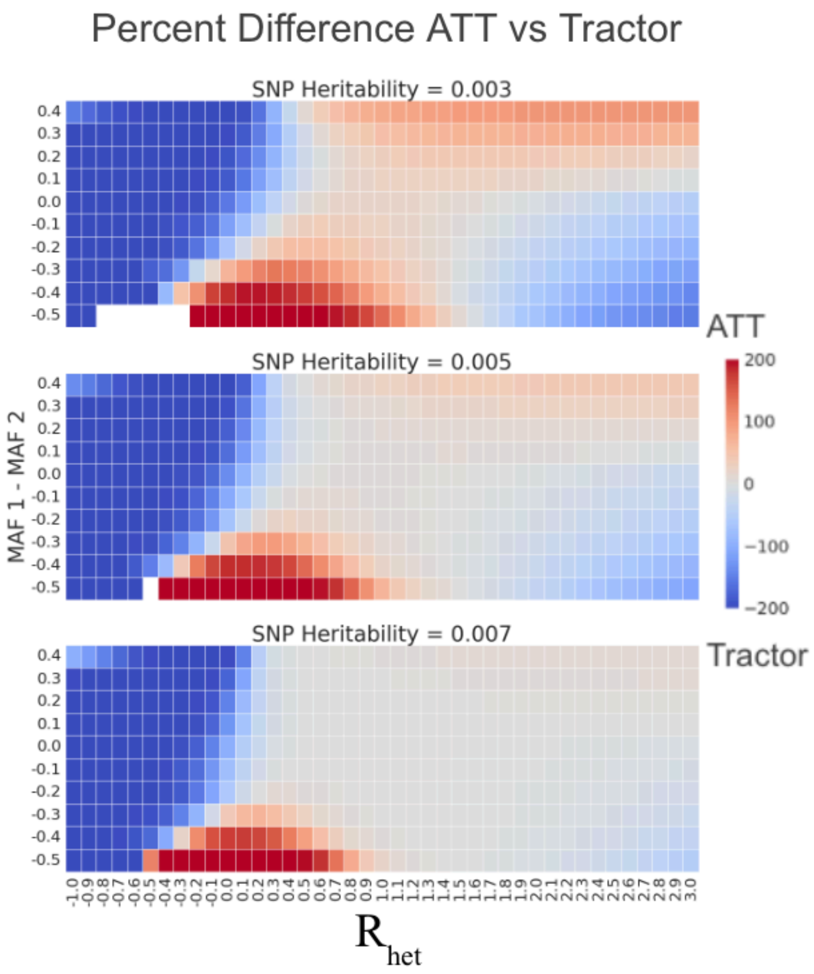


**Figure S7: Impact of HetLanc and MAF difference on percent difference in power depends on heritability.** Heatmap of percent difference in power for ATT vs Tractor. Red indicates where $Power_{ATT} > Power_{Tractor}$. As heritability decreases, the percent difference in power between ATT and Tractor increases. Simulation results of 1,000 replicates with N = 10,000 individuals with minor allele frequency $MAF_1 = 0.5$, global ancestry proportions at 50/50, and causal effect size $\beta_2 = 1.0$.

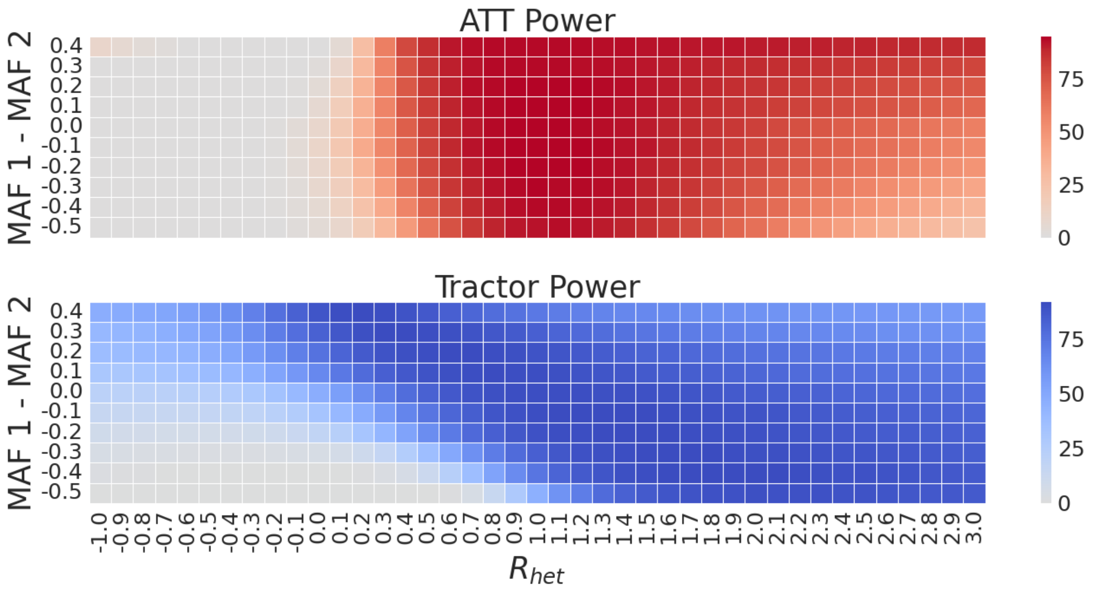


**Figure S8: Impact of HetLanc and MAF difference on power of ATT and Tractor individually.** As HetLanc increases, ATT power decreases, especially when causal effects are in opposite directions. MAF difference impacts Tractor more drastically than ATT. Simulation results of 1,000 replicates with N = 10,000 individuals with minor allele frequency $MAF_1 = 0.5$, global ancestry proportions at 50/50, heritability $h^2 = 0.005$, and causal effect size $\beta_2 = 1.0$.

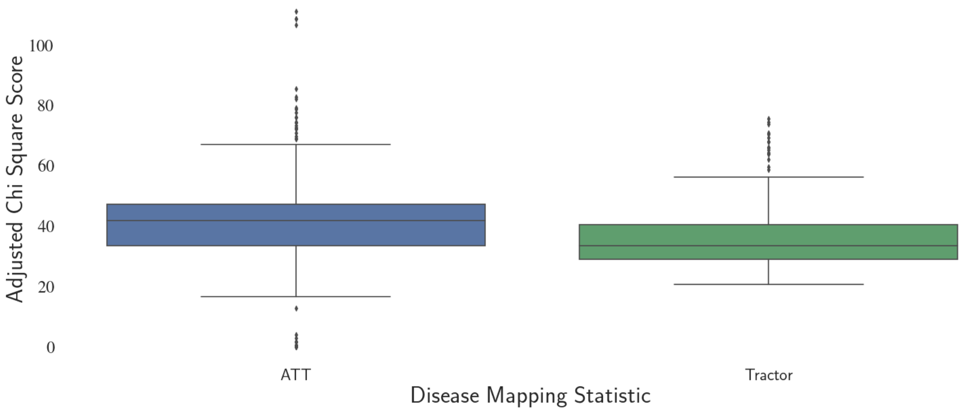


**Figure S9: Adjusted Chi Square Statistics for significant SNPs for 12 traits in the UKBB.** ATT $\chi_1^2$ is significantly larger than the Tractor statistic (adjusted from $\chi_2^2$ to $\chi_1^2$). Mean ATT $\chi_1^2$ for significant SNPs 42.9, mean Tractor $\chi_2^2$ for significant SNPs 37.5, p-value $2.11 \times 10^{-4}$. Study population is 4,327 individuals from the UK Biobank with on average 58.9% African and 41.1% European admixed ancestry. Tractor and ATT statistics computed over 16,584,433 SNPs and 12 traits including AST, BMI, cholesterol, erythrocyte count, HDL, height, LDL, leukocyte count, lymphocyte count, monocyte count, platelet count, and triglycerides. See methods for chi-square adjustment.

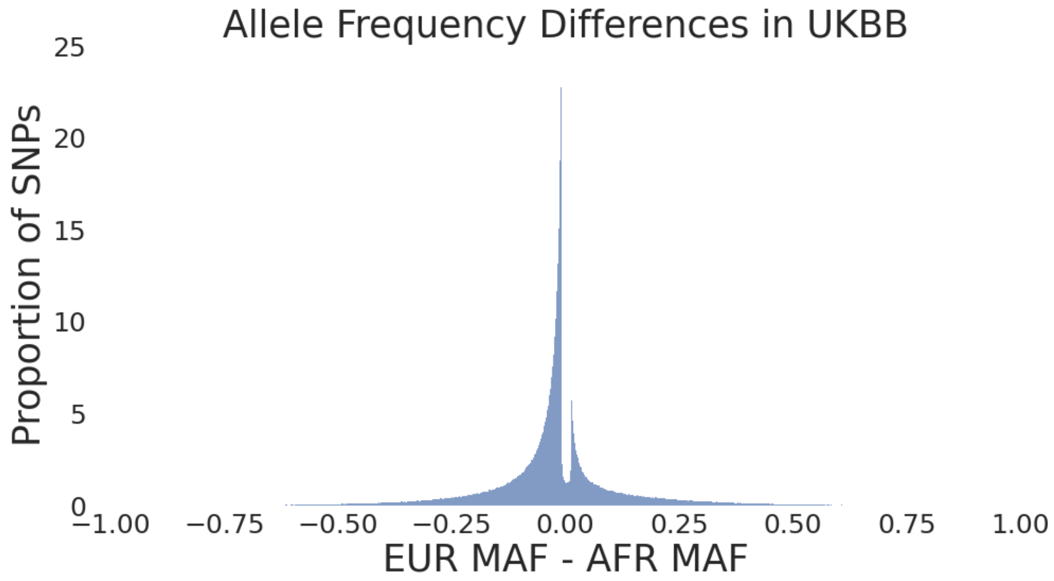


**Figure S10: Minor allele frequency differences in between European and African local ancestries in the African-European admixed population in the UKBB.** Minor allele frequency differences center near zero, at $-2.39 \times 10^{-2}$, indicating only a small systematic bias towards larger minor allele frequencies in the African local ancestry segments. Mean absolute value of minor allele frequency differences is $9.59 \times 10^{-2}$, indicating a small average allele frequency difference, with a standard deviation of $1.15 \times 10^{-1}$. Study population is 4,327 individuals from the UK Biobank with on average 58.9% African and 41.1% European admixed ancestry.

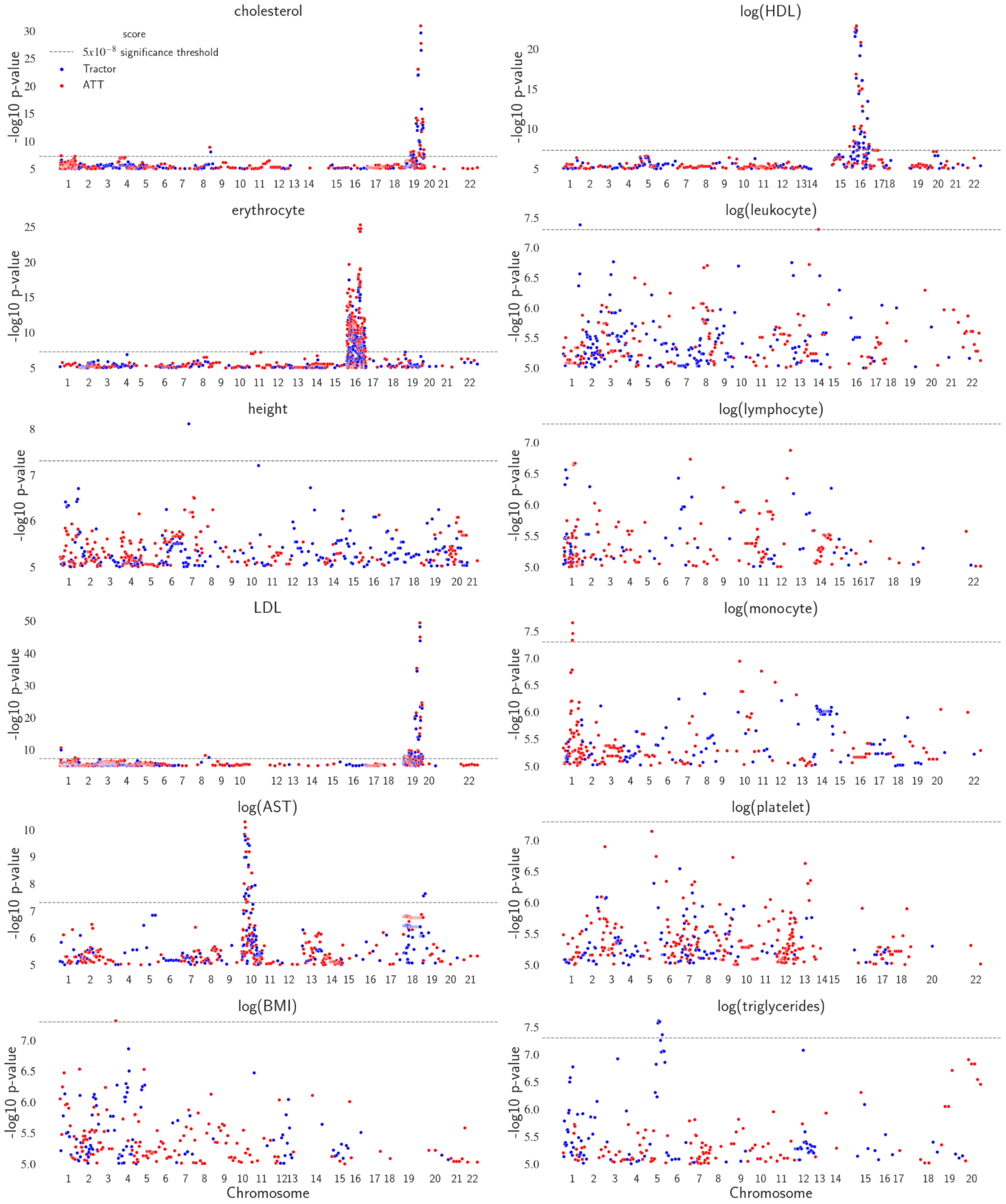


**Figure S11: Manhattan plots for 12 quantitative traits in the UKBB African-European admixed population.** Study population is 4,327 individuals from the UK Biobank with on average 58.9% African and 41.1% European admixed ancestry.

**Table S1: Number of Significant Loci by Phenotype**

| Phenotype | # Loci ATT | # Loci Tractor | # Loci Shared |
|---|---|---|---|
| cholesterol | 3 | 2 | 2 |
| erythrocyte | 3 | 3 | 2 |
| height | 0 | 1 | 0 |
| LDL | 4 | 3 | 3 |
| log(AST) | 1 | 1 | 0 |
| log(BMI) | 1 | 0 | 0 |
| log(HDL) | 5 | 0 | 0 |
| log(leukocyte) | 1 | 0 | 0 |
| log(lymphocyte) | 0 | 0 | 0 |
| log(monocyte) | 1 | 0 | 0 |
| log(platelets) | 0 | 0 | 0 |
| log(triglycerides) | 0 | 0 | 0 |

**Table S2: Significant SNPs in UKBB Admixed Population**

| Phenotype | SNP | ATT p-value | Tractor p-value |
|---|---|---|---|
| cholesterol | chr1:55054772 | $3.72 \times 10^{-8}$ | not significant |
| cholesterol | chr8:118543713 | $1.19 \times 10^{-9}$ | $8.31 \times 10^{-9}$ |
| cholesterol | chr19:44908822 | $1.22 \times 10^{-31}$ | $2.31 \times 10^{-30}$ |
| erythrocyte | chr16:261108 | $5.44 \times 10^{-26}$ | not significant |
| erythrocyte | chr16:360054 | $9.15 \times 10^{-13}$ | not significant |
| erythrocyte | chr16:50884914 | $4.92 \times 10^{-10}$ | not significant |
| erythrocyte | chr16:117409 | not significant | $3.47 \times 10^{-18}$ |
| erythrocyte | chr16:260355 | not significant | $6.34 \times 10^{-18}$ |
| erythrocyte | chr16:384271 | not significant | $2.33 \times 10^{-11}$ |
| height | chr7:78824856 | not significant | $7.79 \times 10^{-9}$ |
| LDL | chr1:55063542 | $2.47 \times 10^{-11}$ | $1.14 \times 10^{-10}$ |
| LDL | chr1:88869866 | $3.01 \times 10^{-8}$ | not significant |
| LDL | chr8:118543713 | $5.74 \times 10^{-9}$ | $2.45 \times 10^{-8}$ |
| LDL | chr19:44908822 | $3.58 \times 10^{-50}$ | $6.24 \times 10^{-49}$ |
| log(AST) | chr10:17819068 | $5.03 \times 10^{-11}$ | not significant |
| log(AST) | chr19:17024164 | not significant | $2.30 \times 10^{-8}$ |
| log(BMI) | chr3:196672134 | $4.83 \times 10^{-8}$ | not significant |

| | | | |
|---|---|---|---|
| log(HDL) | chr15:76063105 | $1.54 \times 10^{-8}$ | not significant |
| log(HDL) | chr16:56957451 | $1.34 \times 10^{-8}$ | not significant |
| log(HDL) | chr17:58519260 | $4.30 \times 10^{-8}$ | not significant |
| log(HDL) | chr17:58607316 | $4.94 \times 10^{-8}$ | not significant |
| log(HDL) | chr17:58744530 | $4.94 \times 10^{-8}$ | not significant |
| log(leukocyte) | chr14:30683993 | $4.96 \times 10^{-8}$ | not significant |
| log(monocyte) | chr1:159092646 | $2.21 \times 10^{-8}$ | not significant |
